# Identification of novel interacting proteins of FUZ and GPR161

**DOI:** 10.1101/2025.09.19.677430

**Authors:** Gabriella Salazar, Anna Battenhouse, Andrew J Kim, Sung-Eun Kim

## Abstract

Proteins modulate signaling pathways through dynamic protein-protein interactions, which provide critical insights into the underlying mechanisms of cellular processes leading to human diseases. Our previous study revealed the biochemical and genetic interactions between FUZ and GPR161 that regulate sonic hedgehog signaling during spinal neural tube development. This study aimed to identify novel co-interacting proteins of FUZ and GPR161 and explore their biochemical and functional associations. We performed affinity-based liquid chromatography–tandem mass spectrometry on immunoprecipitated FUZ and GPR161 proteins. Using this approach, we identified 159 co-interacting proteins, of which 289 proteins that interacted exclusively with FUZ and 617 proteins that interacted exclusively with GPR161. Gene Ontology (GO) analysis of the FUZ and GPR161 co-interactome revealed an enrichment of proteins associated with proteasomal catabolic processes and trafficking. GO analysis of the exclusive FUZ and GPR161 interactomes identified cell cycle progression, mitochondrial membrane, RNA metabolism, receptor complex, and Endoplasmic Reticulum-Golgi transport. These findings were further validated using STRING network analysis. Among the identified proteins, we prioritized FKBP8 and confirmed its biochemical interactions with both FUZ and GPR161 exclusively. In summary, our proteomic profiling uncovered the protein network of FUZ and GPR161, revealing their individual and cooperative functions in multiple cellular processes.

**Statement of Significance:**

- The functional link between FUZ and GPR161 remains unknown, while our previous study showed their genetic and biochemical link in regulating sonic hedgehog signaling during spinal neural tube development.
- This study identified the protein network involving FUZ and GPR161 using affinity- based mass spectrometry combining with gene ontology and STRING analysis, providing the novel molecular regulatory role of FUZ and GPR161.
- We prioritized the proteins, FKBP8, as a co-interacting partners of FUZ and GPR161, and validated their biochemical interaction. We identified the FUZ-GPR161-FKBP8 forms protein complex, and their interaction was not exclusive, suggesting those three proteins located in the same signaling pathway potentially regulating spinal neural tube development.

## • Introduction

Protein-protein interaction (PPI) involves physical and biochemical interactions between proteins in a functional domain-dependent manner. Protein complexes regulated by PPI control multiple cellular processes in a context-dependent manner. Most PPIs are transient and specific in response to extracellular stimuli and the conditions of adjacent proteins. This nature of PPI allows multiple signaling pathways to transduce specific commands to cells for appropriate cellular outcomes. In addition, genetic mutations linked to human diseases inevitably cause protein dysfunction, regardless of whether they are coding or non-coding. One of the landmarks of functional defects in proteins due to genetic mutations is disrupted PPI. For instance, compromised PPI due to genetic modification of candidate genes in neural tube defects (NTDs) has been widely investigated ^[1–3]^ to understand the functional role of individual proteins resulting from mutated genes found in NTD patients. Collectively, a comprehensive PPI network allows us to better understand human diseases.

G-protein-coupled receptor 161 (GPR161) is a G protein-coupled receptor (GPCR), which is the largest receptor family in the human proteome. GPR161 localizes to primary cilia to function as a negative regulator of the sonic hedgehog (Shh) signaling pathway via the Gαs-cAMP-PKA axis ^[4]^. As the ciliary localization of GPR161 is critical for its basal suppressive activity on Shh signaling via regulating Gli2 and Gli3 processing, trafficking in and out of the primary cilia is the key process to determine GPR161 activity. In addition, GPR161 acts as an A-kinase anchored protein (AKAP), thereby facilitating the compartmentalization of GPR161 into the plasma membrane and spatiotemporal PKA recruitment to GPR161 ^[5]^. Therefore, dynamic PPI is required for proper GPR161 activity to regulate Shh signaling pathway. Loss-of-function and dominant negative mutations of *GPR161* in humans and mice are associated with several pediatric diseases, including multiple birth defects, pediatric tumors, and pituitary stalk interruption syndrome ^[6–12]^. However, its associated protein network has not been fully investigated.

The Fuzzy planar cell polarity protein, FUZ, is a Ciliogenesis and PLANar polarity effector (CPLANE) protein complex that is critical for proper ciliogenesis ^[13,14]^. FUZ also facilitates retrograde ciliary trafficking as a retrograde IFT complex ^[15]^. Loss-of-function mutants of *Fuz* in mice present with NTDs, limb abnormalities, and craniofacial defects ^[13,16]^. Mutations in the FUZ gene in humans are linked to NTDs, oral-facial-digital syndrome, and craniosynostosis ^[17–19]^. Considering the role of primary cilia in Shh signaling and the range of phenotypic malformations in *Fuz* mutants in humans and mice, Fuz is also considered a part of Shh signaling ^[20]^, although its role has not been well characterized. Our recent work ^[21]^ revealed a genetic and biochemical link between FUZ and GPR161 in the regulation of Shh signaling during neural tube development. While we showed the evidence that FUZ may be involved in GPR161 trafficking within primary cilia ^[21]^, their biochemical and functional links have not been fully elucidated.

In this study, we identified the proteins that interact with FUZ and GPR161, as well as the proteins that interact with both FUZ and GPR161, using affinity purification-mass spectrometry. Furthermore, we validated the biochemical interactions among FUZ, GPR161, and the prioritized protein. Our results reveal novel protein networks involving both FUZ and GPR161 and provide unexplored functional roles for these two proteins potentially relevant to spinal neural tube development.

## 2. Materials and Methods

### DNA plasmids

The lentiviral vector (*pLV-eGFP*) and packaging vectors (*pRSV-Rev, pMDLg/pRRE, and pMD2.G*) were obtained from Addgene. hGPR161 genes were in-frame inserted into *pLV-eGFP* with BamHI, and *hFUZ* genes were in-frame inserted into *pLV-eGFP* with XbaI and BamHI. The expression vectors for *hGPR161-HA* ^[6]^, *hGPR161-GFP, hFUZ-mCherry* ^[21]^, *and hFKBP8-GFP* ^[1]^ were described in previous publications.

### Cell culture and transfection

HEK293 cells (ATCC-CRL-1573) and HEK293T cells (ATCC-CRL-3216) cells were cultured in Dulbecco’s Modified Eagle Medium (DMEM) with 1% penicillin-streptomycin (PS) and 10% Fetal Bovine Serum (FBS) at 37°C and 5% CO_2_. For overexpression studies, HEK293 and HEK293T cells were transfected with the respective DNA plasmids and Lipofectamine 2000 (Life Technologies) and incubated for 48 h, according to the manufacturer’s instructions.

### Lentivirus production and infection

For lentivirus production, 293T cells were transfected with *pLV-eGFP*, *pLV-GPR161-eGFP*, or *pLV-FUZ-eGFP*, together with *pRSV-Rev*, *pMDLg/pRRE*, and *pMD2.G* using Lipofectamine 2000. The virus-containing medium was collected 48 h post-transfection, and fresh medium was added. The virus-containing medium was collected every 12 h for two to three additional times. All collected virus-containing media were centrifuged at 3000 rpm for 20 min at 4°C. The supernatant was collected and stored at −80°C until infection. For lentivirus infection, 293 cells were plated and infected with virus-containing medium with 8 μg/ml polybrene for 24 h, followed by replacement with fresh media and incubation for an additional 24 h until lysis.

### Co-Immunoprecipitation

HEK293T and HEK293 cells were either transfected with expression vectors indicated or infected with lentivirus containing GFP, GPR161-GFP, or FUZ-GFP and lysed with digitonin-based GPCR immunoprecipitation (GPCR IP) lysis buffer, as previously described ^[22]^. Lysates were immunoprecipitated with ChromoTek GFP-Trap agarose (Proteintech) at 4°C overnight, followed by washing with GPCR IP lysis buffer. For HA IP, lysates were immunoprecipitated with either GFP or HA antibody with protein G agarose at 4°C overnight, followed by washing with GPCR IP lysis buffer. For mass spectrometry, the agarose after IP was washed vigorously (extra washes for extended time) to reduce the non-specific binding. The IP lysates were subjected to SDS-PAGE, followed by western blotting (WB) or in-gel digestion for mass spectrometry.

### Western Blot

Whole cell lysates or immunoprecipitated lysates were subjected to SDS-PAGE, followed by WB. The primary antibodies used were as follows: GFP (Cell signaling #2956; 1:1000), HA (Cell signaling #3724; 1:1000), and RFP (MBL PM005; 1:3000). The secondary antibodies used were IRDye® 800CW goat anti-rabbit IgG and IRDye® 680RD goat anti-mouse IgG (Li-COR). Images were obtained using an Odyssey® scanner (Li-COR).

### Liquid chromatography mass spectrometry (LC-MS/MS)

Immunoprecipitated lysates were subjected to run on the pre-cast 4-20% gradient proteins gels (BioRad) for 10 min. Gels were stained with Coomassie staining solution (Bio-Rad), were excised, and were diced into 1mm gel cubes with ethanol-washed blades for in-gel trypsin digestion, as described by Goodman et al. ^[23]^. Proteins were digested with trypsin, digested peptides were desalted C18 resin-filled pipette tips (Thermo Scientific) following the manufacturer’s instruction, and identified by LC-MS/MS on a Thermo Ultimate 3000 RSLCnano UPLC and Orbitrap Fusion Tribrid mass spectrometer with a 2-hour run time, followed by Sequest HT database search in Proteome Discoverer 2.2 and Scaffold validation in the UT Austin Biological Mass Spectrometry Facility using previously published method ^[24,25]^.

### Database search

LC-MS/MS raw files were obtained for protein A agarose (two biological replicates) GFP (three biological replicates), FUZ-GFP (three biological replicates), GPR161-GFP (three biological replicates) IP samples and were converted to mzXML format with MSconvert ^[26]^ using standard parameters, then analyzed using X!Tandem ^[27]^, MSGF+ ^[28]^ and Comet ^[29]^ to match mass spectra to peptide sequences. All data were searched against the human UniProt ^[30]^ proteome, UP000005640. All searches employed a 10-ppm mass tolerance for both precursor and fragment ions for higher energy collisional dissociation (HCD). Enzyme specificity was set fully specific to trypsin (cleavage sites were RK, cleavage side was C-terminal) with up to two missed cleavages allowed. Oxidation of methionine was used as a common modification. Peptide matches from the three search engines were probabilistically integrated using MSblender ^[31]^ and assigned to proteins as spectral counts. Data were filtered to a 1% false discovery rate (FDR) using decoy searching with common contaminants added.

### Bait/prey interaction analysis

The SAINTexpress (v3.6.3) ^[32]^ software package was used to discriminate between bona fide protein-protein interactions in affinity purified mass spectrometry experiments from nonspecific background interactions as modeled by control purifications. SAINTexpress input files (bait, prey and interactions) were formatted according to recommendations in the original SAINT analysis protocol document ^[33]^. Specifically: total spectral counts were used for quantification; spectral counts for the bait itself were removed from its own experiments; and the control experiments (GFP and protein A) were treated as separate baits rather than replicates of a control type. The saintexp-spc runs were performed with the -R 2 command line option (use the 2 best-scoring interactions when more than 2 replicate observations are available). ^[32]^ The outputs from the SAINTexpress runs were combined and annotated. Significance levels of baits were determined with two parameters:at least 1 unique peptide spectral match (PSM) and SAINT Baysian false discovery rate (FDR) <=0.05. Two significance subsets were further annotated: Significance 1 (Sig 1) refers to at least one unique PSM and FDR ≤ 0.01, and significance 2 (Sig 2) refers to at least one unique PSM and FDR ≤ 0.05. Co-interacting proteins of FUZ and GPR161 were selected from baits with Sig 1 and Sig 2 from both FUZ and GPR161.

### Functional enrichment analysis

Gene Ontology (GO) analysis was performed in R (v3.6.1) using the topGO package (v2.38.1). Sig 1 and Sig 2 interacting proteins were first converted to corresponding gene IDs using GENCODE v34 ^[34]^ human gene annotations. Three data subsets (FUZ-only, GPR161-only, and FUZ+GPR161) were analyzed separately for the 3 GO ontologies (Biological Process/BP, Molecular Function/MF and Cellular Component/CC) against the org.Hs.eg.db annotation database using topGO’s classic algorithm and Fisher’s Exact test in count data mode. Protein network analysis based on known functional enrichment was performed using web-based STRING analysis (https://string-db.org, version 12.0).

## 3. Results

Our previous study ^[21]^ revealed that *Fuz* and *Gpr161* are genetically linked to the regulation of Shh signaling, spinal neural tube development, and their biochemical interactions. Since GPR161 is mainly localized in the primary cilia ^[4]^, while FUZ is known to localize in the basal body to regulate ciliogenesis ^[13]^, we speculate that their biochemical interaction may not be a direct protein-protein interaction. Hence, we sought to identify the protein interactomes of FUZ and GPR161 to characterize their molecular functions.

To explore the biochemical and functional links between FUZ and GPR161, we sought to identify the protein complexes that interact with FUZ and GPR161 using proteomic approaches. To this end, we performed affinity-based liquid chromatography with tandem mass spectrometry (LC-MS/MS) analysis using FUZ and GPR161. The overall experimental scheme is shown in Figure 1, while the basic validation experiments are shown in Supplementary Figure 1. In brief, 293 cells were infected with lentivirus containing GFP, FUZ-GFP, or GPR161-GFP. Cell lysates were immunoprecipitated with GFP nanobody, which is the smallest antibody with high affinity and low non-specific binding, to minimize non-specific binding to the IgG antibody structure. In addition, we included vigorous controls, including two protein A bead-only control pull down with cells overexpressing either FUZ-GFP or GPR161-GFP, and three GFP Trap with cells overexpressing GFP only, to rule out the possibility of immunoprecipitation (IP) artifacts and overexpression and protein tag artifacts. We selected HEK293 cells, which are human ciliated cells ^[35]^, because the localization of GPR161 and FUZ is relevant to primary cilia.

**Figure 1.**
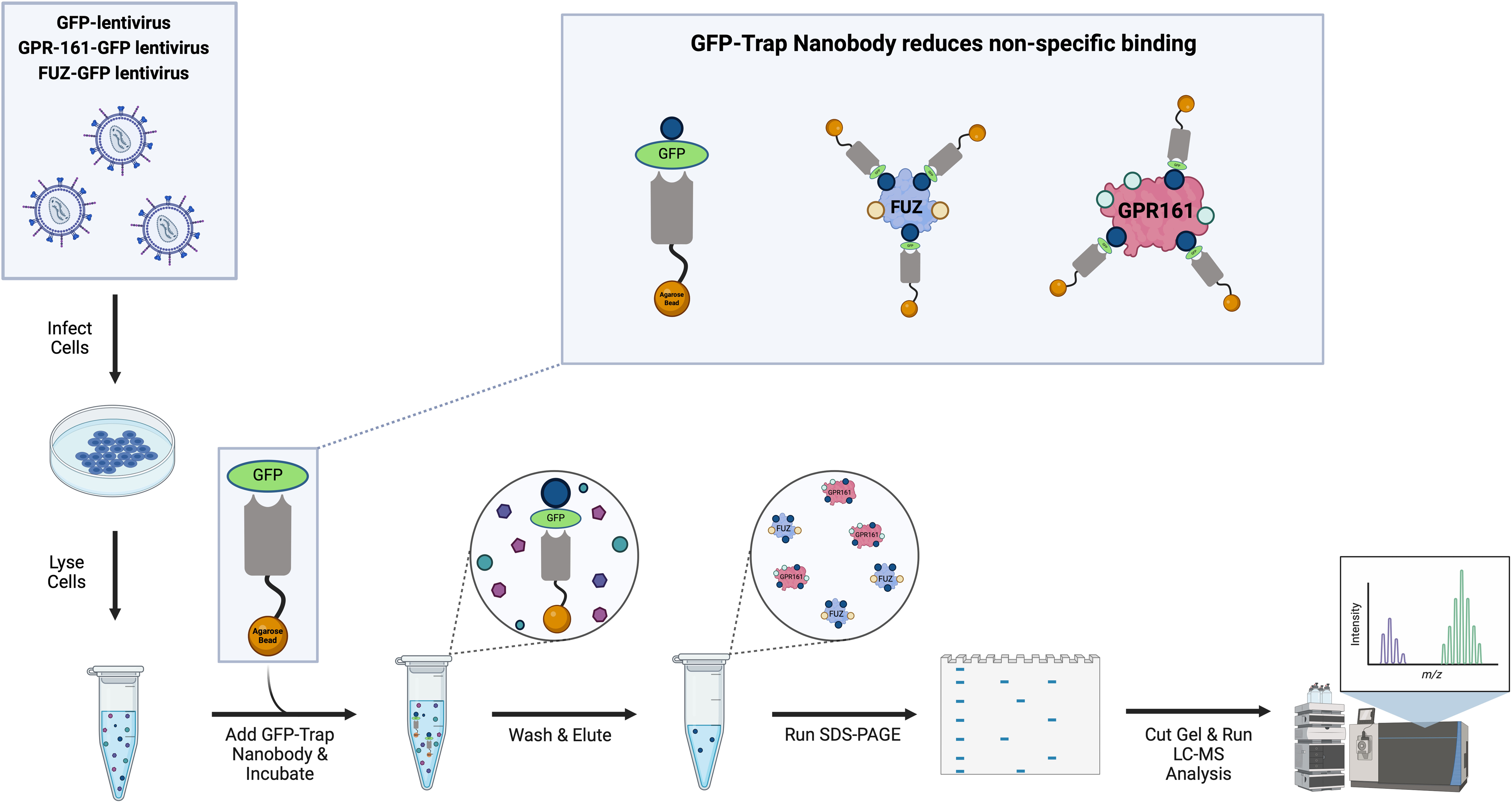
Overview of affinity purification-based LC-MS/MS experiments. 293T cells were infected with lentivirus containing GFP, FUZ-GFP, or GPR161-GFP before lysis. The lysates were subjected to immunoprecipitation with GFP nanobody-conjugated agarose to pull down the protein complex associated with GFP, FUZ-GFP, or GPR161-GFP proteins. After vigorous washing, the protein complexes with the antibody-agarose complex were denatured, run on SDS-PAGE, and cut for in-gel digestion for mass spectrometry.

The LC-MS/MS data were searched using the human UniProt database, and the peptide spectra counts were assigned to each protein. We identified proteins that significantly interacted with either FUZ or GPR161 using the Significance Analysis of INTeractome software (SAINTexpress analysis ^[32]^). To identify the co-interacting proteins for FUZ and GPR161, we sorted the proteins that included both FUZ and GPR161 interactome with the highest significance based on SAINTexpress analysis. We identified a total of 776 proteins that significantly interacted with GPR161 and 448 proteins that interacted with FUZ. Among them, 159 proteins were identified to significantly interact with both GPR161 and FUZ (Figure 2A).

**Figure 2.**
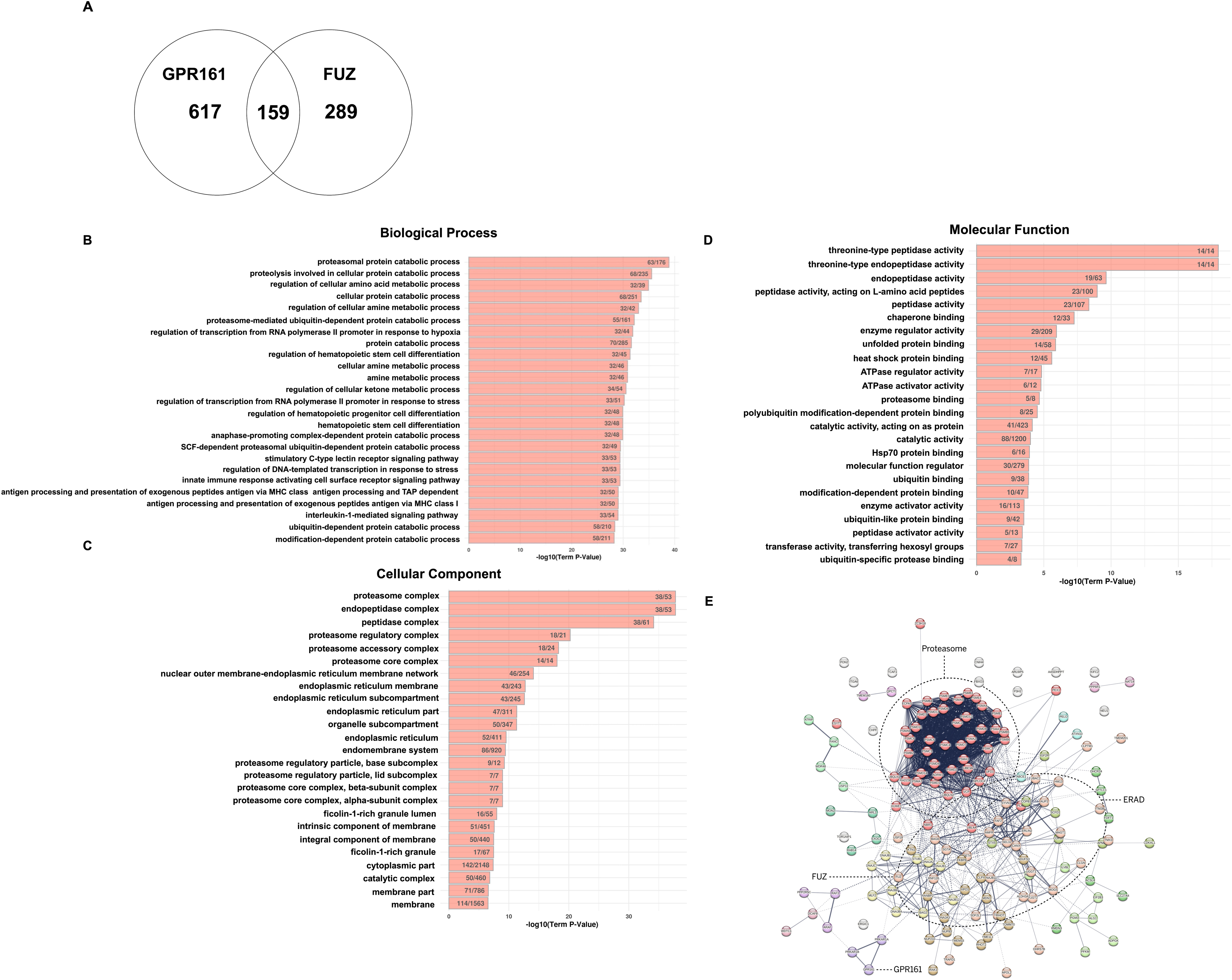
Identification and analysis of the co-interactome of FUZ and GPR161. (A) Venn diagram indicating the number of identified proteins that interact with both FUZ and GPR161 and exclusively interact with either FUZ or GPR161. (B-D) Gene Ontology analysis of proteins that interact with both FUZ and GPR161. Top 25 protein categories for Biological Process (B), Cell Component (C), and Molecular Function ontologies (D). The numbers on each bar graph represent the number of proteins identified within each GO term category, relative to the total number of proteins in that category. (E) STRING analysis of proteins interacting with both FUZ and GPR161. Proteins in the same category are color-coded. Two main nodes, the proteasome and ERAD, are indicated by circles.

### Analysis co-interactome between FUZ and GPR161

We first performed Gene Ontology (GO) analysis to classify the FUZ and GPR161 co-interactome to determine their functional enrichment (Figure 2B-2D). The top five biological processes in the co-interactome of FUZ and GPR161 included proteasomal protein catabolic pathways, proteolysis involved in the cellular protein catabolic process, regulation of the cellular amino acid metabolic process, cellular protein catabolic process, and regulation of the cellular amine metabolic process (Figure 2B). The top five cellular components were the proteasome complex, endopeptidase complex, peptidase complex, proteasome regulatory particle, and proteasome core complex (Figure 2C). The top five molecular functions were threonine-type peptidase activity, threonine-type endopeptidase activity, L-amino acid peptidase activity, peptidase activity, and chaperone binding (Figure 2D). Intriguingly, the top terms in all three ontologies in the GO analysis of the FUZ–GPR161 co-interactome were involved in functional processes relevant to protein degradation. To explore the functional association between individual proteins within the FUZ–GPR161 co-interactome, we performed STRING analysis ^[36]^. The proteins within the co-interactome were functionally connected based on the existing protein interaction database in the STRING analysis (Figure 2E). Two large nodes within this connectome included the proteasome complex and ER-associated degradation (ERAD) proteins (Figure 2E). There were 48 proteins in the proteasome complex and 31 proteins in the ERAD category, as shown in Supplementary Figure 2. The notable overlap between the GO term analysis and our table reinforces the central role of the co-interactome in protein processing and degradation. We then categorized 159 individual proteins into proteasomes, chaperones, ubiquitin-dependent ligases and related proteins, trafficking proteins, enzymes relevant to post-translational modification, and uncategorized proteins based on the protein functions described in the Uniprot protein database (Supplementary Table).

### Analysis of FUZ interactome

We examined proteins that exclusively interact with FUZ. As shown in Figure 1B, we identified 448 proteins that significantly interacted with FUZ. Among them, INTU, RSG, and OFD1 were included, which are known FUZ-interacting proteins. We also compared the published FUZ interactome ^[14]^ to the FUZ interactome we identified, and there were 162 proteins present in both interactomes (Supplementary Figure 3). We further examined 287 proteins that exclusively interacted with FUZ. Based on GO analysis, the top five biological processes included proteins enriched in mitotic cell cycle regulation and post-transcriptional gene silencing (Figure 3A), which is consistent with their role in ciliogenesis. Proteins were enriched in the ubiquitin ligase complex and nucleoplasm in the top five cellular components and regulatory transcription factors, including repressors or binding, in the top five molecular functions (Figure 3A). We further examined the proteins that exclusively interact with FUZ using STRING analysis (Figure 3B). There were three large nodes within the FUZ interactome: RNA and nucleic acid metabolic processes (57 proteins), mitosis (55 proteins), and mitochondrial membrane organization (31 proteins) (Supplementary Figure 4). Notably, mitotic proteins were annotated using both GO and STRING analyses.

**Figure 3.**
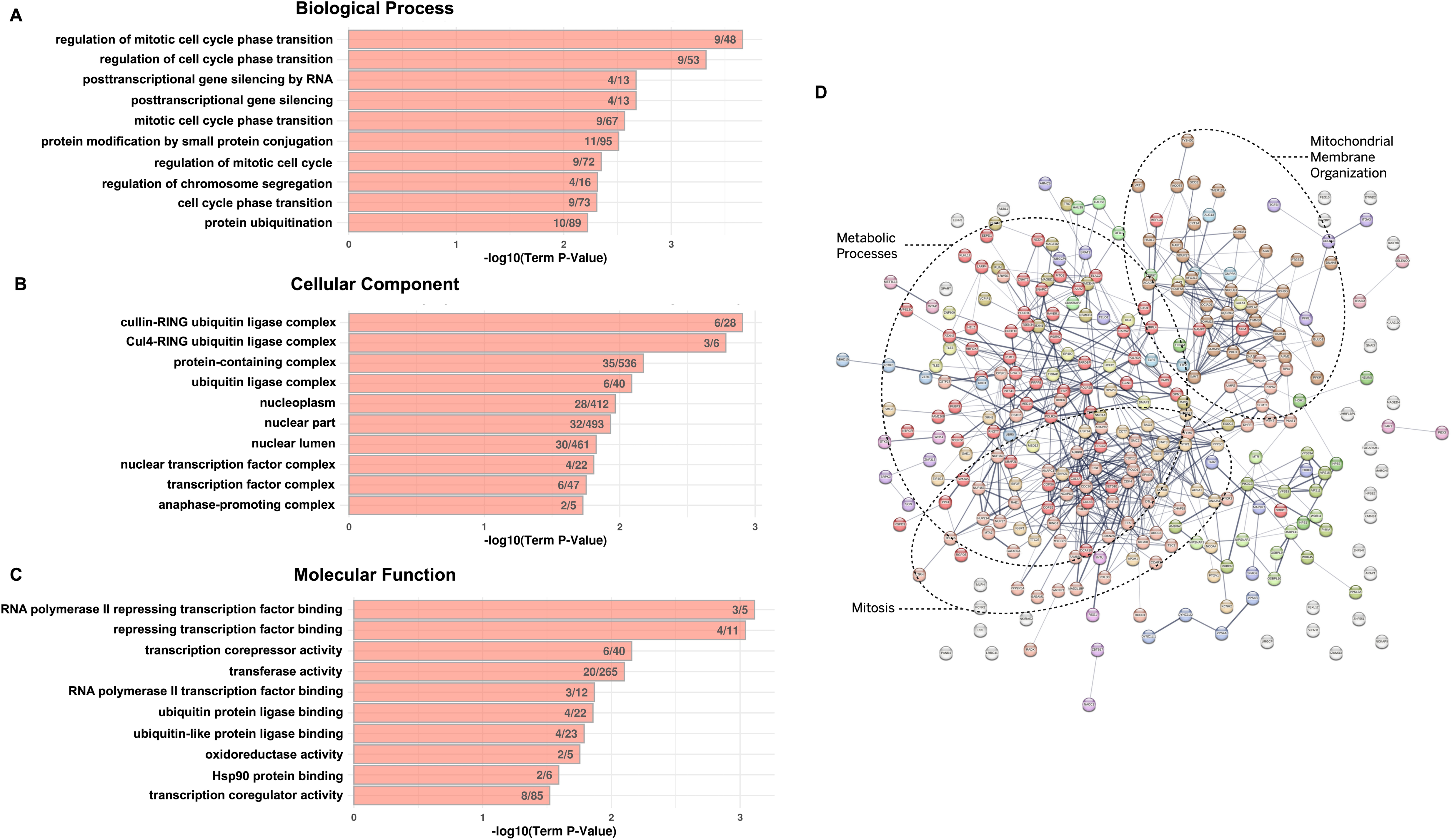
Identification and analysis of the exclusive interactome of FUZ. (A-C) Gene Ontology analysis of FUZ-exclusive interacting proteins. Top 10 protein categories for Biological Process (A), Cell Component (B), and Molecular Function ontologies (C). The numbers on each bar graph represent the number of proteins identified within each GO term category, relative to the total number of proteins in that category. (D) STRING analysis of FUZ-exclusive interacting proteins. Proteins in the same category are color-coded. Three main nodes, metabolic process, mitosis, and mitochondrial membrane organization are indicated as circles.

### Analysis of GPR161 interactome

We analyzed the GPR161 interactome and found that PRKAR1A, PRKAR1B, AKAP4, and multiple PKA catalytic subunits (PRKACA, PRKACB, PRKACG, and PRKACSH) have been previously identified as GPR161 interacting proteins in the literatures ^[5,37]^. Among the GPR161 interactome, we identified 617 proteins that exclusively interacted with GPR161 (Figure 2A). Based on GO analysis, proteins were enriched in localization, glycoprotein metabolic processes, and transport in the Biological Process (Figure 4A). Proteins were enriched in membrane components in the top five cellular components (Figure 4B) and in transmembrane receptor protein kinase, molecular transducer, signaling receptor, and UDP-glycosyltransferase activity in the top five molecular functions (Figure 4C). The GO results support the well-known role of GPR161 as a GPCR. We then performed STRING analysis using the proteins that interacted exclusively with GPR161 (Figure 4D). Two main nodes were identified in the GPR161 interactome, including the enzyme-linked receptor protein signaling pathway (136 proteins) and ER-to-Golgi vesicle-mediated transport (71 proteins) (Supplementary Figure 5). Enzyme-linked receptor protein signaling pathways were identified by both STRING and GO analyses, suggesting the specific receptor function of GPR161.

**Figure 4.**
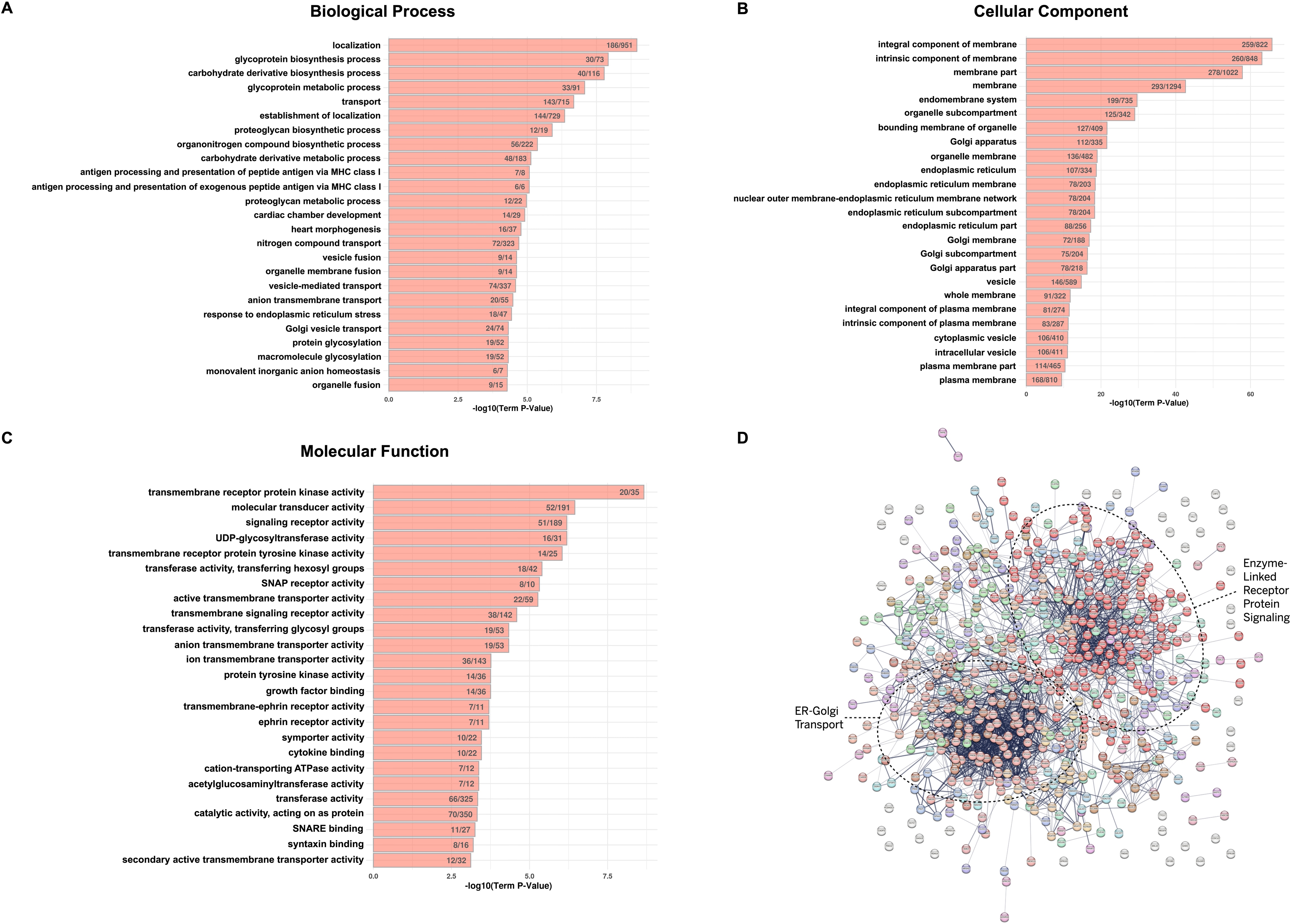
Identification and analysis of the exclusive interactome of GPR161. (A-C) Gene Ontology analysis of GPR161 exclusive interacting proteins. Top 25 protein categories for Biological Process (A), Cell Component (B), and Molecular Function ontologies (C). The numbers on each bar graph represent the number of proteins identified within each GO term category, relative to the total number of proteins in that category. (D) STRING analysis of GPR161 exclusive interacting proteins. Proteins in the same category are color-coded. Two main nodes, metabolic process, enzyme-linked receptor protein signaling, and ER-Golgi transport, are indicated as circles.

### Validation of the co-interactome: FKBP8 is a co-interacting protein of FUZ and GPR161

We prioritized the co-interactome of FUZ and GPR161 using the following three criteria for the validation study. First, it is related to neural tube development; second, it is relevant to primary cilia; and finally, it is a Shh signaling molecule, as both FUZ and GPR161 fall into all three criteria. FKBP8 satisfied all three criteria among co-interacting proteins of FUZ and GPR161, and FKBP8 was also identified as FUZ interacting proteins in the previous study ^[14]^. First, we validated the biochemical interaction between GPR161 and FKBP8 by performing co-immunoprecipitation experiments with overexpressed GPR161 and FKBP8 proteins in HEK293 cells. FKPB8 and GPR161 proteins were immunoprecipitated using GPR161 pulldown (Figure 5A) and FKBP8 pulldown (Figure 5B), respectively, confirming their biochemical interaction. We then examined the interactions between all three proteins, FKBP8, FUZ, and GPR161. We co-overexpressed all three genes, FUZ, GPR161, and FKBP8, and then immunoprecipitated with GPR161. The GPR161-FKBP8 biochemical interaction was not affected by FUZ overexpression, whereas all three proteins were in the protein complex (Figure 5C). In addition, we further confirmed the individual biochemical interactions between GPR161 and FUZ (Figure 5D: lanes 1 and 2), and between FKBP8 and FUZ (Figure 5D: lanes 3 and 4). Similarly, the biochemical interaction between FKBP8 and FUZ was not changed by GPR161 co-expression, while FKBP8 still interacts with GPR161 (Figure 5D: compare between lane 4 and 5). In addition, we predicted the structural features of these interactions using Alpha fold 3 ^[38]^. Consistent with our protein interaction data in Figure 5D, structural interaction interfaces among full-length of FKBP8, FUZ and GPR161 were not overlapping (Supplementary Figure 6). However, these predictions were considered to be a low confidence interaction (ipTM: 0.18 and pTM: 0.46 for FUZ and GPR161, ipTM: 0.24 and pTM: 0.41). Taken together, these results validate our proteomics data showing FKBP8 interacts with both FUZ and GPR161 and further suggest that the biochemical interaction among three proteins is not exclusive.

**Figure 5.**
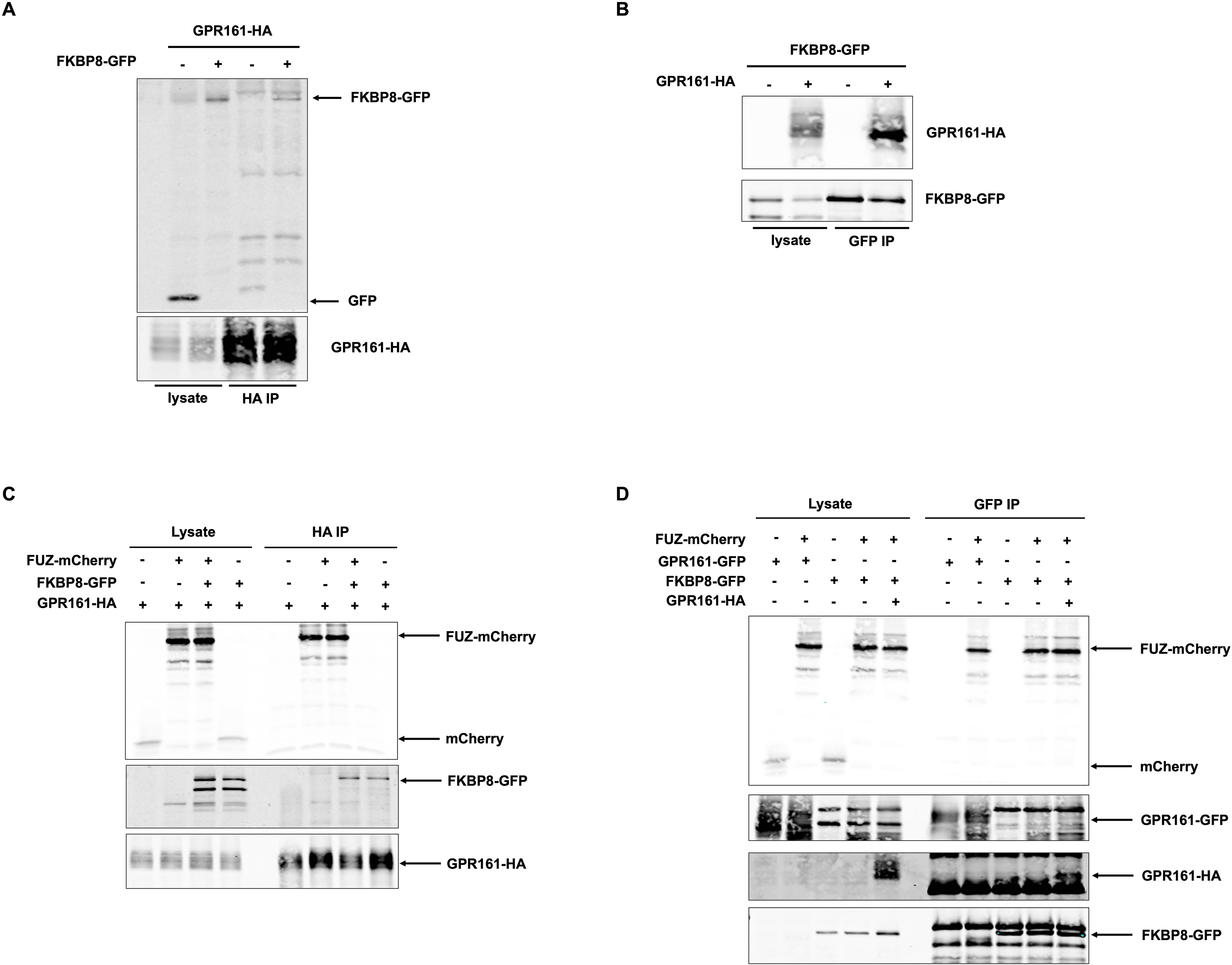
Validation study with a prioritized protein: FKBP8. (A-B) Validation of the biochemical interaction between GPR161 and FKBP8. 293 cells were transfected with FKBP8-GFP and GPR161-HA plasmids, and cell lysates were subjected to immunoprecipitation (IP) with an anti-HA antibody, followed by western blotting (WB) to detect GPR161-HA and FKBP8-GFP (A). IP was performed using an anti-GFP nano-Trap, followed by WB to detect GPR161-HA and FKBP8-GFP (B). (C and D) Biochemical interactions between FUZ, GPR161, and FKBP8. 293 cells were transfected with GPR161-HA, FKBP8-GFP, and FUZ-mCherry plasmids, and the cell lysates were subjected to IP with anti-HA antibody (C) or anti-GFP antibody (D), followed by WB to detect GPR161-HA GPR161-GFP, FKBP8-GFP, and FUZ-mCherry. (A-D) Representative data from n=2 for each IP experiment. IP: Immunoprecipitation.

## 4. Discussion

In this study, we identified the protein interactome of FUZ and GPR161 to understand their functional connection using affinity-based mass spectrometry. We then categorized the identified interacting proteins using GO and STRING analyses and determined the novel functions of both FUZ and GPR161 and individual FUZ and GPR161. Finally, we prioritized the protein as an interacting partner of both FUZ and GPR161, potentially implying this protein complex relevant to either the regulation of Shh signaling or the spinal neural tube development.

Two notable protein categories in the co-interactome of FUZ and GPR161 are protein catabolism and trafficking. Within protein catabolism, we observed both the proteasome complex and ERAD proteins and E3-ligases as two large nodes from the co-interactome (Figure 2), suggesting that FUZ and GPR161 can be regulated by ubiquitin-dependent degradation for proteostasis regulation. We ruled out the possibility that it could be an overexpression artifact and non-specific binding with abundant proteins for two reasons: 1) we removed the interacting proteins of the GFP control from the FUZ and GPR161 interactomes, which used the same overexpression system and affinity purification methods, and 2) milder overexpression with lentivirus infection. Remarkably, GPR161 is known to be ubiquitinated at Lys63 for its removal from cilia mediated by the BBSome, with the coupling of endosomal sorting required for transport (ESCRT)-0 protein ^[39]^. This work is aligned with our previous finding that FUZ, as one of the retrograde intraflagellar (IFT) machinery, might be involved in β-arrestin-mediated ciliary removal of GPR161 ^[21]^. A recent publication further supports that both BBSome and IFT machinery mediate b-arrestin-dependent ciliary removal of GPR161 ^[40]^. While the ubiquitination of GPR161 at Lys63 is linked to BBSome-dependent GPR161 ciliary removal, the involvement of IFT in this process remains unclear. Our study provides evidence that FUZ, an IFT machinery component, may serve to remove ubiquitin-tagged GPR161 from cilia. In addition, the proteomics data that we identified further provide information regarding specific ubiquitin ligases and the potential link between ciliary removal of ubiquitinated GPR161 and its degradation, both of which could be mediated by FUZ. This will be a compelling follow-up study.

Among the trafficking categories, ciliary trafficking is a particular area of interest given that FUZ and GPR161 are both known to have critical roles within primary cilia. We identified 18 proteins related to ciliary trafficking, including ARL1, ARL6IP5, BAG6, DNAJB6, EXOC4, FKBP8, IGF2R, KIFC2, KTN1, NUP210, NUP93, PEX1, PRKAR1A, PRKAR1B, PSMD2, RAB14, UGGT1, and VCP, of 32 proteins categorized to trafficking proteins (Supplementary Table). Among them, FKBP8 was of particular interest since FKBP8 has a role in Shh signaling and an implication in cilia (both motile and non-motile) ^[41,42]^. Furthermore, genetic mutations in Fkbp8 are risk factors for spina bifida in humans and Fkbp8 knockout mice have 100% penetrant spina bifida ^[1,43]^. Taken together, these factors are all consistent with the functional and genetic roles of FUZ and GPR161. Considering the known role of FKBP8, it may be involved in ciliogenesis together with FUZ or act as a co-chaperone of both FUZ and GPR161 for protein homeostasis. According to our validation IP experiment (Figure 5), their interaction is not exclusive, suggesting that their binding domains may not be overlapping within each protein. In particular, the functional relevance of either regulation in the context of spinal neural tube development will be the focus of our future study.

The exclusive FUZ interactome suggests the potential novel functions of FUZ during mitosis. Taken into account its localization in the basal body of primary cilia, the basal body is a remnant of centriole, which plays a critical role in organizing the mitotic spindle of the chromosome during mitosis. Our data (Figure 3) supports the notion that FUZ interacts with proteins related to mitosis, which might lead to its functional role during cell cycle progression. In addition, ubiquitin ligase complex/binding was identified as a FUZ interactome by GO analysis (Figure 3A-3C), suggesting that FUZ may be regulated by ubiquitin-dependent degradation machinery. This novel protein regulatory mechanism of FUZ has implications for birth defects, including NTDs, considering that the loss-of-function mutations of FUZ are associated with NTDs and ciliopathies ^[17,44,45]^.

The exclusive GPR161 interactomes included two major categories of proteins: enzyme-linked receptor protein signaling and ER-Golgi transport (Figure 4). Considering the role of GPR161 as a receptor of Shh signaling, it is reasonable to identify multiple other receptors for signaling. These results suggest that GPR161 may form protein complexes with other receptors in other signaling pathways, thereby potentially regulating other signaling pathways. Indeed, our previous studies ^[6–8]^ suggest that GPR161 regulates Shh and Wnt signaling, although the molecular regulatory mechanisms have not been fully elucidated. These data provide potential mechanistic insights into our observations. The other remarkable nodes of the STRING analysis with the GPR161 exclusive interactome were ER-Golgi transport. While GPCR is well known to transport from ER to Golgi, then to plasma membrane, our result showed the specific protein complex for ER to Golgi transport for GPR161. In addition, Popa et al. ^[46]^ revealed that GPR161 is a Golgi-resident protein that maintains the Golgi structure. Our results are also aligned with this notion, suggesting that GPR161 is not only transported via ER-Golgi transporters but also regulates Golgi homeostasis for its proper secretion and that of other proteins.

Overall, our proteomics study identified the protein network of FUZ and GPR161, which sheds light on the novel functions of FUZ and GPR161, either together or individually. Despite the limitations of overexpression systems, our study provides molecular clues to improve the scientific understanding of the molecular functions of FUZ and GPR161 and to identify novel diagnostic markers relevant to the diseases in which FUZ and GPR161 are involved.

## 5. Associated Data

Mass spectrometry raw and processed data was submitted to MassIVE (MSV000098847), and will be publicly available upon publication.

## Acknowledgement

In-gel digestion, mass spectrometry, and initial protein identification were provided by Michelle Gadush and Maria Person in the UT Austin Center for Biomedical Research Support (CBRS) Biological Mass Spectrometry Facility (RRID: SCR_021728). Computational analyses were performed by the Biomedical Research Computing Facility at UT Austin, CBRS (RRID: SCR_021979). We thank Cyndi Liang for initial alpha fold analysis in Supplementary Figure 6 and Dr. Richard Finnell (Baylor College of Medicine) for the critical comments. Schematic image in Figure 1 was generated using BioRender, and the manuscript was proofread with Paperpal.

## Author contribution

SEK conceived the overall study; GS and SEK performed the overall experiments; and AB, GS, and AJK analyzed the proteomics dataset; GS and SEK drafted the manuscript, and all authors reviewed and edited the manuscript.

## References

[1] Tian, T., Cao, X., Kim, S. E., Lin, Y. L., Steele, J. W., Cabrera, R. M., … Lei, Y. (2020). FKBP8 variants are risk factors for spina bifida. Hum Mol Genet, 29(18), 3132–3144. doi: 10.1093/hmg/ddaa211

[2] Lei, Y., Fathe, K., McCartney, D., Zhu, H., Yang, W., Ross, M. E., … Finnell, R. H. (2015). Rare LRP6 variants identified in spina bifida patients. Hum Mutat, 36(3), 342–349. doi: 10.1002/humu.22750

[3] Lei, Y., Kim, S. E., Chen, Z., Cao, X., Zhu, H., Yang, W., … Finnell, R. H. (2019). Variants identified in PTK7 associated with neural tube defects. Mol Genet Genomic Med, 7(4), e00584. doi: 10.1002/mgg3.584

[4] Mukhopadhyay, S., Wen, X., Ratti, N., Loktev, A., Rangell, L., Scales, S. J., & Jackson, P. K. (2013). The ciliary G-protein-coupled receptor Gpr161 negatively regulates the Sonic hedgehog pathway via cAMP signaling. Cell, 152(1-2), 210–223. doi: 10.1016/j.cell.2012.12.026

[5] Bachmann, V. A., Mayrhofer, J. E., Ilouz, R., Tschaikner, P., Raffeiner, P., Rock, R., … Stefan, E. (2016). Gpr161 anchoring of PKA consolidates GPCR and cAMP signaling. Proc Natl Acad Sci U S A, 113(28), 7786–7791. doi: 10.1073/pnas.1608061113

[6] Kim, S. E., Lei, Y., Hwang, S. H., Wlodarczyk, B. J., Mukhopadhyay, S., Shaw, G. M., … Finnell, R. H. (2019). Dominant negative GPR161 rare variants are risk factors of human spina bifida. Hum Mol Genet, 28(2), 200–208. doi: 10.1093/hmg/ddy339

[7] Kim, S. E., Robles-Lopez, K., Cao, X., Liu, K., Chothani, P. J., Bhavani, N., … Finnell, R. H. (2021). Wnt1 Lineage Specific Deletion of Gpr161 Results in Embryonic Midbrain Malformation and Failure of Craniofacial Skeletal Development. Front Genet, 12, 761418. doi: 10.3389/fgene.2021.761418

[8] Kim, S. E., Chothani, P. J., Shaik, R., Pollard, W., & Finnell, R. H. (2023). Pax3 lineage-specific deletion of Gpr161 is associated with spinal neural tube and craniofacial malformations during embryonic development. Dis Model Mech, 16(11). doi: 10.1242/dmm.050277

[9] Hwang, S. H., White, K. A., Somatilaka, B. N., Shelton, J. M., Richardson, J. A., & Mukhopadhyay, S. (2018). The G protein-coupled receptor Gpr161 regulates forelimb formation, limb patterning and skeletal morphogenesis in a primary cilium-dependent manner. Development, 145(1). doi: 10.1242/dev.154054

[10] Shimada, I. S., Somatilaka, B. N., Hwang, S. H., Anderson, A. G., Shelton, J. M., Rajaram, V., … Mukhopadhyay, S. (2019). Derepression of sonic hedgehog signaling upon Gpr161 deletion unravels forebrain and ventricular abnormalities. Dev Biol, 450(1), 47–62. doi: 10.1016/j.ydbio.2019.03.011

[11] Karaca, E., Buyukkaya, R., Pehlivan, D., Charng, W. L., Yaykasli, K. O., Bayram, Y., … Lupski, J. R. (2015). Whole-exome sequencing identifies homozygous GPR161 mutation in a family with pituitary stalk interruption syndrome. J Clin Endocrinol Metab, 100(1), E140–147. doi: 10.1210/jc.2014-1984

[12] Begemann, M., Waszak, S. M., Robinson, G. W., Jager, N., Sharma, T., Knopp, C., … Kurth, I. (2020). Germline GPR161 Mutations Predispose to Pediatric Medulloblastoma. J Clin Oncol, 38(1), 43–50. doi: 10.1200/JCO.19.00577

[13] Gray, R. S., Abitua, P. B., Wlodarczyk, B. J., Szabo-Rogers, H. L., Blanchard, O., Lee, I., … Finnell, R. H. (2009). The planar cell polarity effector Fuz is essential for targeted membrane trafficking, ciliogenesis and mouse embryonic development. Nat Cell Biol, 11(10), 1225–1232. doi: 10.1038/ncb1966

[14] Toriyama, M., Lee, C., Taylor, S. P., Duran, I., Cohn, D. H., Bruel, A. L., … Wallingford, J. B. (2016). The ciliopathy-associated CPLANE proteins direct basal body recruitment of intraflagellar transport machinery. Nat Genet, 48(6), 648–656. doi: 10.1038/ng.3558

[15] Brooks, E. R., & Wallingford, J. B. (2012). Control of vertebrate intraflagellar transport by the planar cell polarity effector Fuz. J Cell Biol, 198(1), 37–45. doi: 10.1083/jcb.201204072

[16] Zhang, Z., Wlodarczyk, B. J., Niederreither, K., Venugopalan, S., Florez, S., Finnell, R. H., & Amendt, B. A. (2011). Fuz regulates craniofacial development through tissue specific responses to signaling factors. PLoS One, 6(9), e24608. doi: 10.1371/journal.pone.0024608

[17] Seo, J. H., Zilber, Y., Babayeva, S., Liu, J., Kyriakopoulos, P., De Marco, P., … Torban, E. (2011). Mutations in the planar cell polarity gene, Fuzzy, are associated with neural tube defects in humans. Hum Mol Genet, 20(22), 4324–4333. doi: 10.1093/hmg/ddr359

[18] Barrell, W. B., Adel Al-Lami, H., Goos, J. A. C., Swagemakers, S. M. A., van Dooren, M., Torban, E., … Liu, K. J. (2022). Identification of a novel variant of the ciliopathic gene FUZZY associated with craniosynostosis. Eur J Hum Genet, 30(3), 282–290. doi: 10.1038/s41431-021-00988-6

[19] Singh, S., Nampoothiri, S., Narayanan, D. L., Chaudhry, C., Salvankar, S., & Girisha, K. M. (2024). Biallelic loss of function variants in FUZ result in an orofaciodigital syndrome. Eur J Hum Genet, 32(8), 1022–1026. doi: 10.1038/s41431-024-01619-6

[20] Heydeck, W., Zeng, H., & Liu, A. (2009). Planar cell polarity effector gene*Fuzzy*regulates cilia formation and Hedgehog signal transduction in mouse. Developmental Dynamics, 238(12), 3035–3042. doi: 10.1002/dvdy.22130

[21] Kim, S. E., Kim, H. Y., Wlodarczyk, B. J., & Finnell, R. H. (2024). Linkage between Fuz and Gpr161 genes regulates sonic hedgehog signaling during mouse neural tube development. Development, 151(19). doi: 10.1242/dev.202705

[22] Pal, K., Badgandi, H., & Mukhopadhyay, S. (2015). Studying G protein-coupled receptors: immunoblotting, immunoprecipitation, phosphorylation, surface labeling, and cross-linking protocols. Methods Cell Biol, 127, 303–322. doi: 10.1016/bs.mcb.2014.12.003

[23] Goodman, J. K., Zampronio, C. G., Jones, A. M. E., & Hernandez-Fernaud, J. R. (2018). Updates of the In-Gel Digestion Method for Protein Analysis by Mass Spectrometry. Proteomics, 18(23), e1800236. doi: 10.1002/pmic.201800236

[24] Gao, G., Hausmann, S., Flores, N. M., Benitez, A. M., Shen, J., Yang, X., … Bedford, M. T. (2023). The NFIB/CARM1 partnership is a driver in preclinical models of small cell lung cancer. Nat Commun, 14(1), 363. doi: 10.1038/s41467-023-35864-y

[25] Gadush, M. V., Sautto, G. A., Chandrasekaran, H., Bensussan, A., Ross, T. M., Ippolito, G. C., & Person, M. D. (2022). Template-Assisted De Novo Sequencing of SARS-CoV-2 and Influenza Monoclonal Antibodies by Mass Spectrometry. J Proteome Res, 21(7), 1616–1627. doi: 10.1021/acs.jproteome.1c00913

[26] Chambers, M. C., Maclean, B., Burke, R., Amodei, D., Ruderman, D. L., Neumann, S., … Mallick, P. (2012). A cross-platform toolkit for mass spectrometry and proteomics. Nat Biotechnol, 30(10), 918–920. doi: 10.1038/nbt.2377

[27] Craig, R., & Beavis, R. C. (2004). TANDEM: matching proteins with tandem mass spectra. Bioinformatics, 20(9), 1466–1467. doi: 10.1093/bioinformatics/bth092

[28] Kim, S., & Pevzner, P. A. (2014). MS-GF+ makes progress towards a universal database search tool for proteomics. Nat Commun, 5, 5277. doi: 10.1038/ncomms6277

[29] Eng, J. K., Jahan, T. A., & Hoopmann, M. R. (2013). Comet: an open-source MS/MS sequence database search tool. Proteomics, 13(1), 22–24. doi: 10.1002/pmic.201200439

[30] UniProt, C. (2025). UniProt: the Universal Protein Knowledgebase in 2025. Nucleic Acids Res, 53(D1), D609–D617. doi: 10.1093/nar/gkae1010

[31] Kwon, T., Choi, H., Vogel, C., Nesvizhskii, A. I., & Marcotte, E. M. (2011). MSblender: A probabilistic approach for integrating peptide identifications from multiple database search engines. J Proteome Res, 10(7), 2949–2958. doi: 10.1021/pr2002116

[32] Teo, G., Liu, G., Zhang, J., Nesvizhskii, A. I., Gingras, A. C., & Choi, H. (2014). SAINTexpress: improvements and additional features in Significance Analysis of INTeractome software. J Proteomics, 100, 37–43. doi: 10.1016/j.jprot.2013.10.023

[33] Choi, H., Liu, G., Mellacheruvu, D., Tyers, M., Gingras, A. C., & Nesvizhskii, A. I. (2012). Analyzing protein-protein interactions from affinity purification-mass spectrometry data with SAINT. *Curr Protoc Bioinformatics*, Chapter 8, 8 15 11–18 15 23. doi: 10.1002/0471250953.bi0815s39

[34] Frankish, A., Diekhans, M., Ferreira, A. M., Johnson, R., Jungreis, I., Loveland, J., … Flicek, P. (2019). GENCODE reference annotation for the human and mouse genomes. Nucleic Acids Res, 47(D1), D766–D773. doi: 10.1093/nar/gky955

[35] Lancaster, M. A., Schroth, J., & Gleeson, J. G. (2011). Subcellular spatial regulation of canonical Wnt signalling at the primary cilium. Nat Cell Biol, 13(6), 700–707. doi: 10.1038/ncb2259

[36] Szklarczyk, D., Kirsch, R., Koutrouli, M., Nastou, K., Mehryary, F., Hachilif, R., … von Mering, C. (2023). The STRING database in 2023: protein-protein association networks and functional enrichment analyses for any sequenced genome of interest. Nucleic Acids Res, 51(D1), D638–D646. doi: 10.1093/nar/gkac1000

[37] Hoppe, N., Harrison, S., Hwang, S. H., Chen, Z., Karelina, M., Deshpande, I., … Manglik, A. (2024). GPR161 structure uncovers the redundant role of sterol-regulated ciliary cAMP signaling in the Hedgehog pathway. Nat Struct Mol Biol, 31(4), 667–677. doi: 10.1038/s41594-024-01223-8

[38] Abramson, J., Adler, J., Dunger, J., Evans, R., Green, T., Pritzel, A., … Jumper, J. M. (2024). Accurate structure prediction of biomolecular interactions with AlphaFold 3. Nature, 630(8016), 493–500. doi: 10.1038/s41586-024-07487-w

[39] Shinde, S. R., Mick, D. U., Aoki, E., Rodrigues, R. B., Gygi, S. P., & Nachury, M. V. (2023). The ancestral ESCRT protein TOM1L2 selects ubiquitinated cargoes for retrieval from cilia. Dev Cell, 58(8), 677–693 e679. doi: 10.1016/j.devcel.2023.03.003

[40] Fujii, T., Murai, N., Aso, S., Takatsu, H., Shin, H. W., Katoh, Y., & Nakayama, K. (2025). beta-Arrestin mediates the export of ciliary GPR161 but not Smoothened together with the BBSome and intraflagellar transport machinery. J Cell Sci, 138(20). doi: 10.1242/jcs.263793

[41] Cho, A., Ko, H. W., & Eggenschwiler, J. T. (2008). FKBP8 cell-autonomously controls neural tube patterning through a Gli2- and Kif3a-dependent mechanism. Dev Biol, 321(1), 27–39. doi: 10.1016/j.ydbio.2008.05.558

[42] Mali, G. R., Yeyati, P. L., Mizuno, S., Dodd, D. O., Tennant, P. A., Keighren, M. A., … Mill, P. (2018). ZMYND10 functions in a chaperone relay during axonemal dynein assembly. Elife, 7. doi: 10.7554/eLife.34389

[43] Wong, R. L., Wlodarczyk, B. J., Min, K. S., Scott, M. L., Kartiko, S., Yu, W., … Finnell, R. H. (2008). Mouse Fkbp8 activity is required to inhibit cell death and establish dorso-ventral patterning in the posterior neural tube. Hum Mol Genet, 17(4), 587–601. doi: 10.1093/hmg/ddm333

[44] Tabler, J. M., Barrell, W. B., Szabo-Rogers, H. L., Healy, C., Yeung, Y., Perdiguero, E. G., … Liu, K. J. (2013). Fuz mutant mice reveal shared mechanisms between ciliopathies and FGF-related syndromes. Dev Cell, 25(6), 623–635. doi: 10.1016/j.devcel.2013.05.021

[45] Tabler, J. M., Rice, C. P., Liu, K. J., & Wallingford, J. B. (2016). A novel ciliopathic skull defect arising from excess neural crest. Dev Biol, 417(1), 4–10. doi: 10.1016/j.ydbio.2016.07.001

[46] Popa, S., Villeneuve, J., Stewart, S., Perez Garcia, E., Petrunkina Harrison, A., & Moreau, K. (2019). Genome-wide CRISPR screening identifies new regulators of glycoprotein secretion. Wellcome Open Res, 4, 119. doi: 10.12688/wellcomeopenres.15232.2

