## Supplementary data and table for "Identification of novel interacting proteins of FUZ and GPR161"

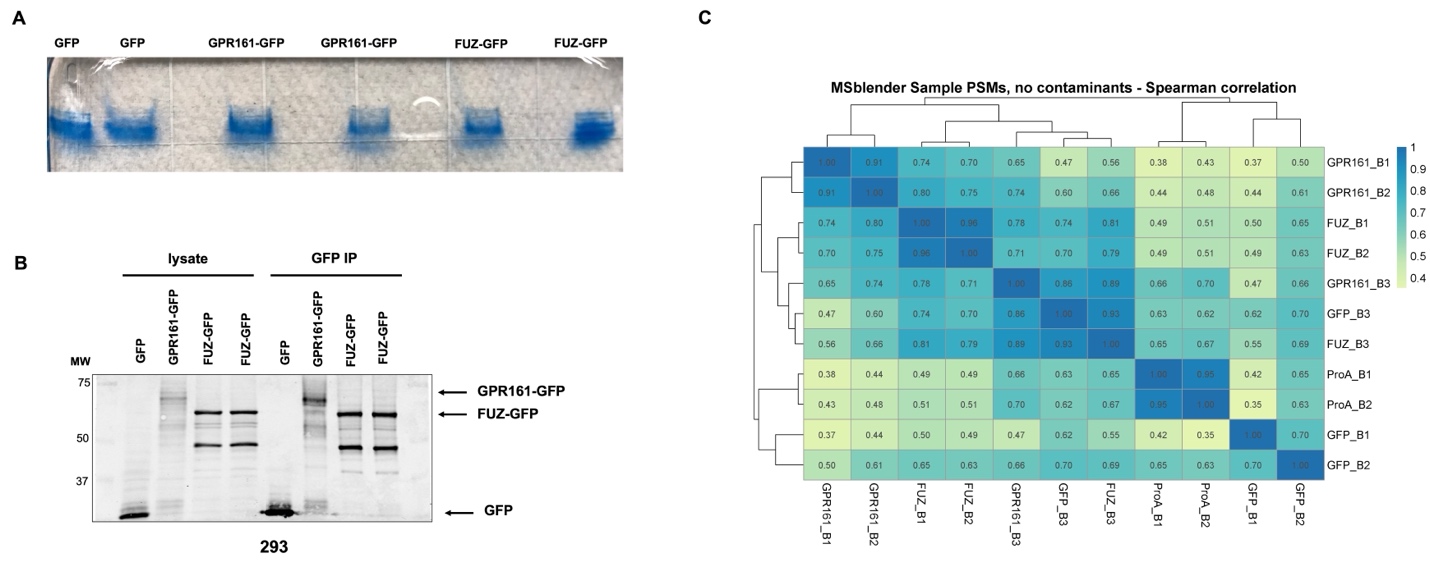

**Supplementary Figure 1. FUZ and GPR161 pull down by IP using GFP nanobody conjugated agarose (GFP nano-Trap).** (A) Coomassie staining gel with two representative GFP, FUZ-GFP, and GPR161-GFP IP samples. Cell lysates from HEK293 cells infected by GFP, FUZ-GFP, and GPR161-GFP lentivirus were pulldown with GFP nanobody conjugated agarose, run SDS-PAGE and subjected to Coomassie staining for obtaining all the prey proteins bound to each bait protein (GFP, FUZ-GFP and GPR161-GFP). (B) IP validation. Small amount of GFP pull down samples from lysates infected with GFP, FUZ-GFP, and GPR161-GFP lentivirus was subjected to WB to detect with anti-GFP antibody. MW: molecular weight. (C) Spearman correlation analysis using PSMs from each pull down sample. 2 protein A pull down samples, 3 GFP pull down samples with GFP infected HEK293 cell lysates, 3 GFP pull down samples with FUZ-GFP infected HEK293 cell lysates, and GFP pull down samples with GPR161-GFP infected HEK293 cell lysates. PSM: peptide spectra matches.

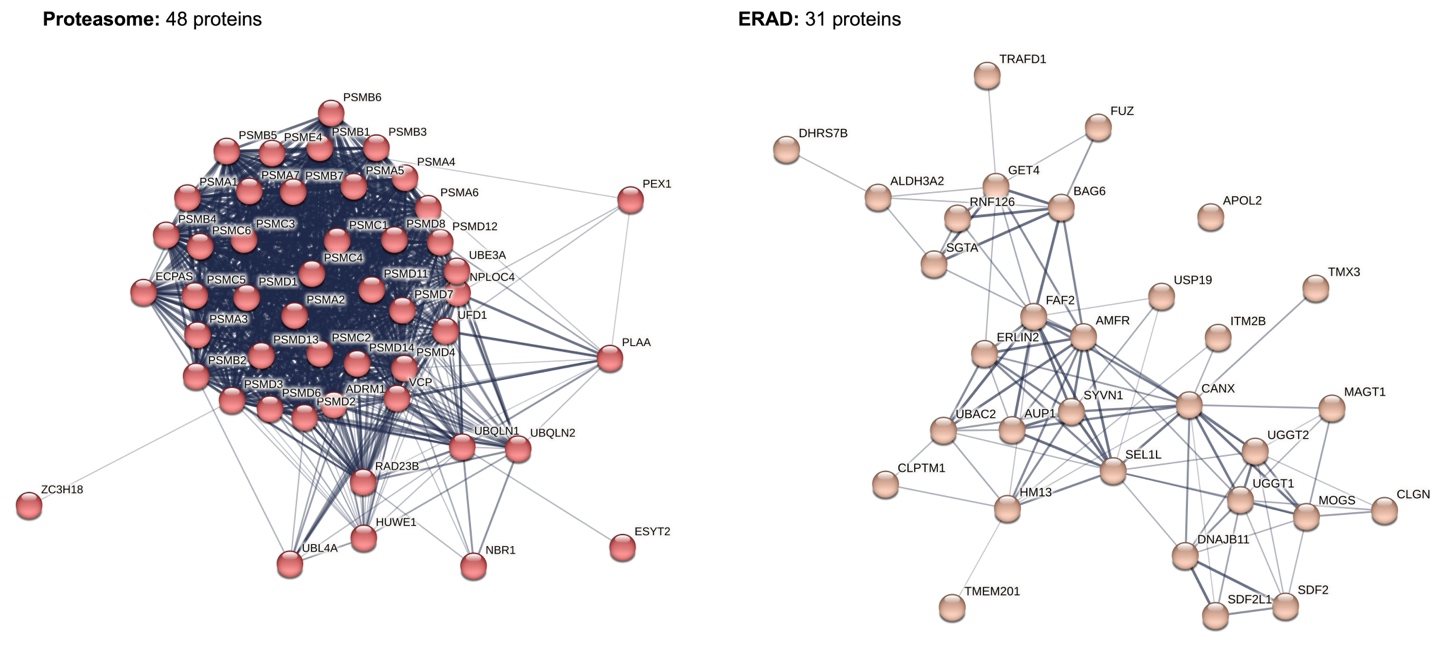

**Supplementary Figure 2.** STRING analysis for two major protein networks of FUZ and GPR161 co-interactome. Two major nodes of protein networks were identified from STRING analysis with FUZ and GPR161 co-interactome. Left: Proteasome and Right: ERAD. The numbers and the names of individual proteins in each protein network were indicated.

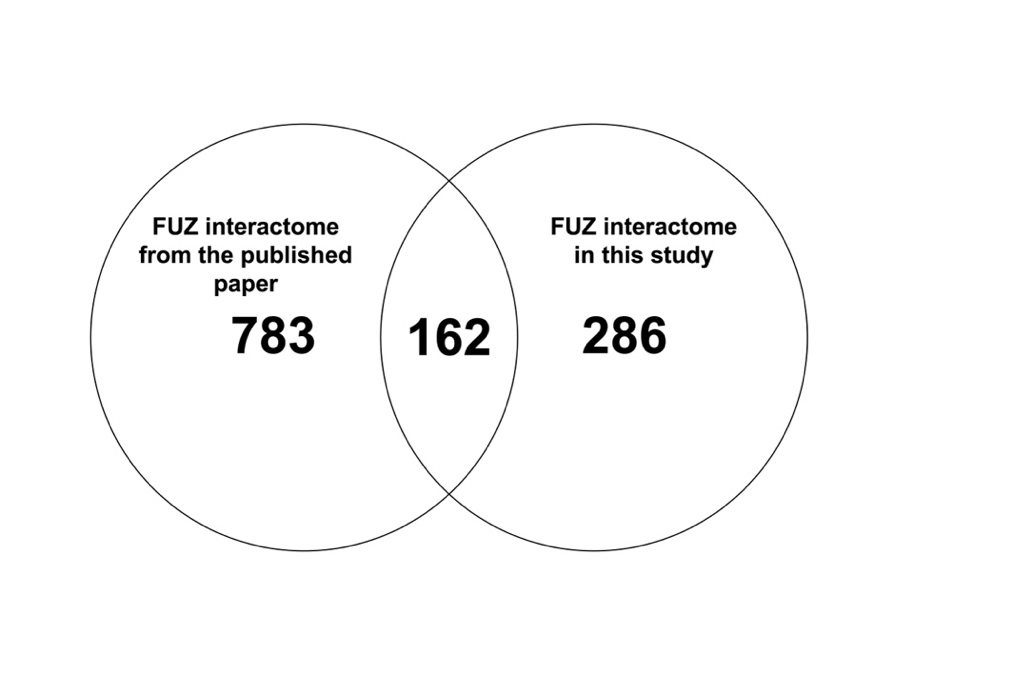

**Supplementary Figure 3.** The comparison of FUZ interactomes between from published work and from this study. We compared two FUZ interactomes and identified 162 proteins shared from two distinct data set. Of note, FUZ interactome from the published work (Ref) was isolated from IMCD3 (mouse) cells and the one from this study was isolated from HEK293 cells (human).

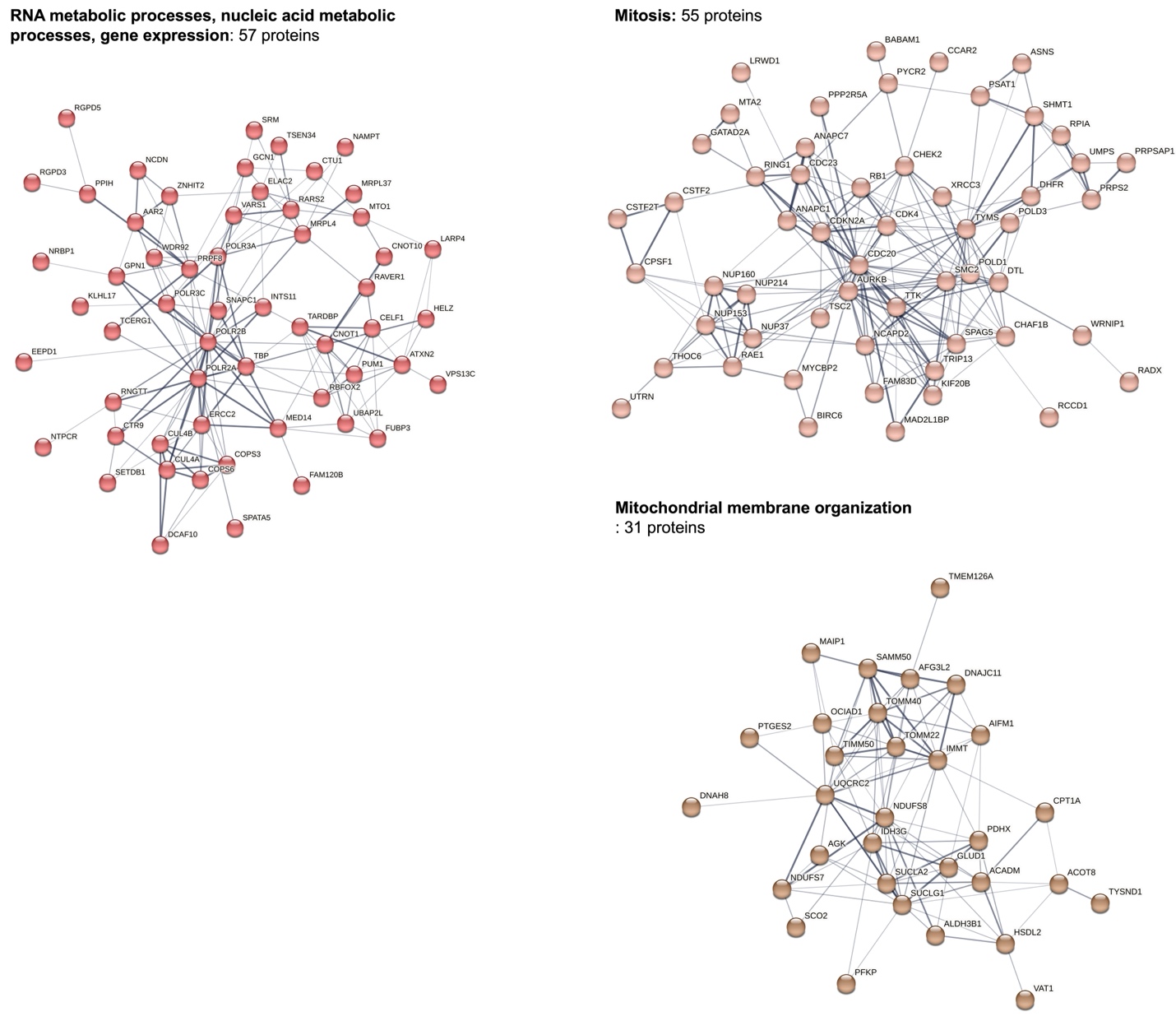

**Supplementary Figure 4.** STRING analysis of three major protein networks from FUZ exclusive interactome. RNA metabolic pathway, mitosis and mitochondria membrane organization. The numbers and the names of individual proteins in each protein network were indicated.

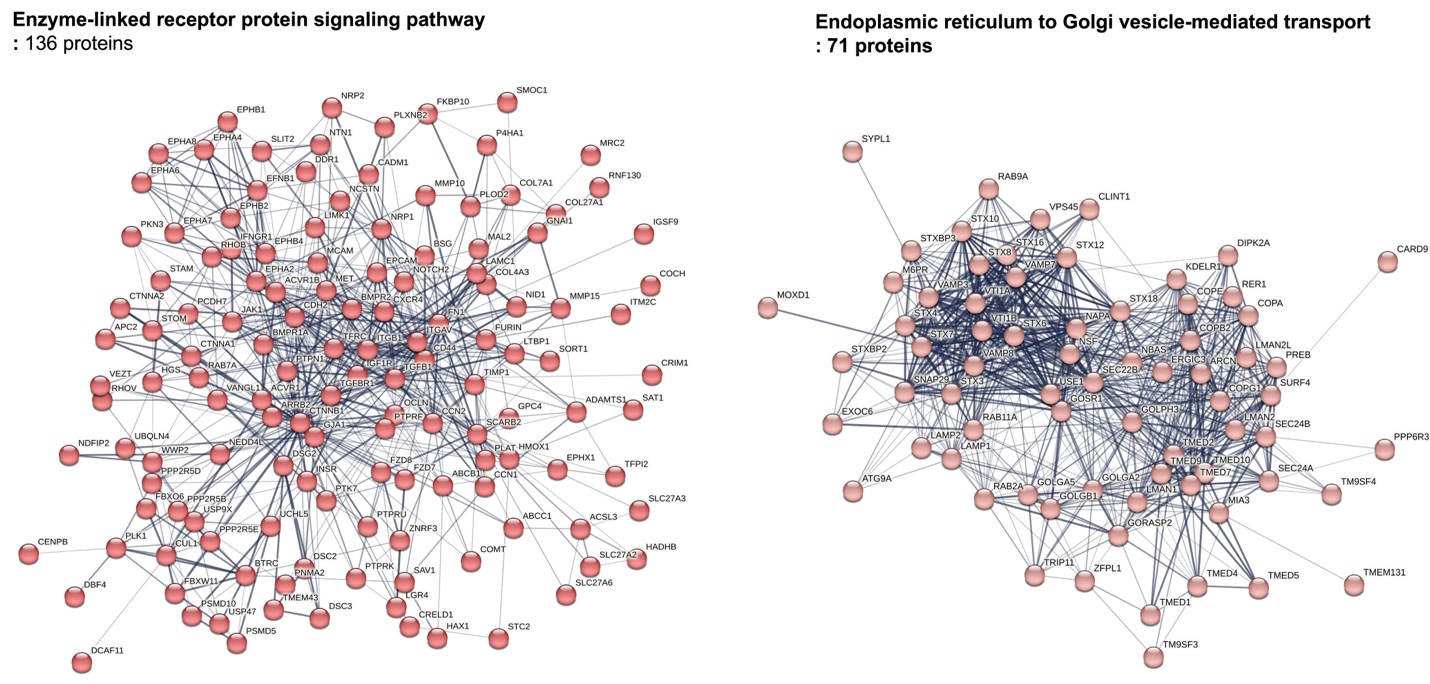

**Supplementary Figure 5.** STRING analysis of two major protein networks from GPR161 exclusive interactome. Enzyme-linked receptor protein signaling pathway and ER-Golgi vesicle-mediated transport. The numbers and the names of individual proteins in each protein network were indicated.

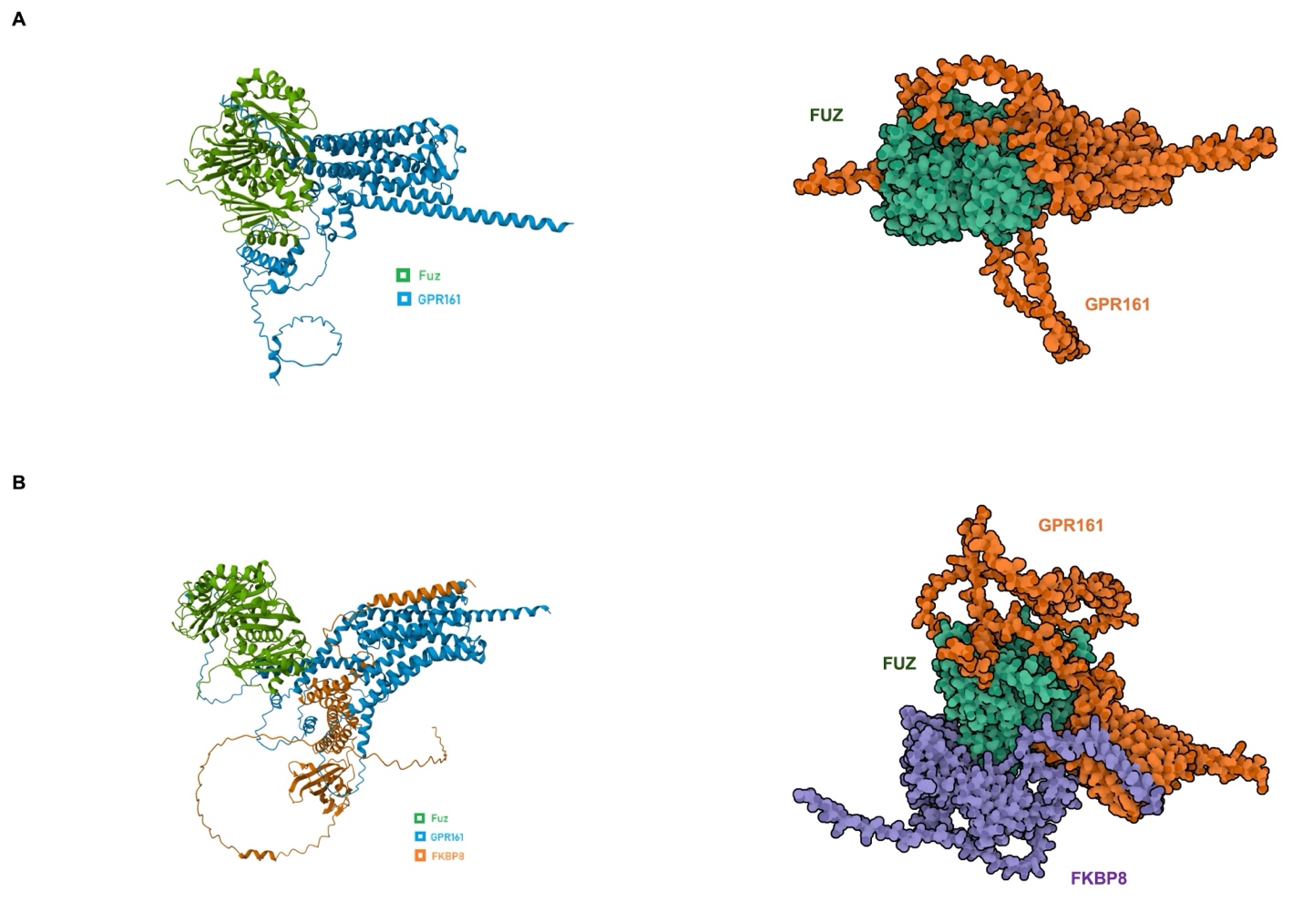

**Supplementary Figure 6.** (A and B) Alpha fold was used to predict the protein-protein interaction between FUZ and GPR16 (A) and among FKBP8, FUZ and GPR161 (B). Left panel: protein complex structures with secondary structures. Right panel: Protein complex structures without secondary structures. Each protein was color-coded.

**Supplementary Table. Categorized protein list with proteins identified from both FUZ and GPR161 interactome.**

| **Name** | **ID** | **Description** |
| --- | --- | --- |
| **Proteasome** | | |
| ADRM1 | sp\|Q16186\|ADRM1_HUMAN | component of 26S proteasome |
| ECM29 | sp\|Q5VYK3\|ECM29_HUMAN | scaffolding protein that binds to the 26S proteasome |
| PSMA1 | sp\|P25786\|PSA1_HUMAN | component of the 20S core proteasome; proteolytic degradation |
| PSMA2 | sp\|P25787\|PSA2_HUMAN | component of the 20S core proteasome; proteolytic degradation |
| PSMA3 | sp\|P25788\|PSA3_HUMAN | component of the 20S core proteasome; proteolytic degradation |
| PSMA4 | sp\|P25789\|PSA4_HUMAN | component of 26S proteasome |
| PSMA5 | sp\|P28066\|PSA5_HUMAN | component of the 20S core proteasome; proteolytic degradation |
| PSMA6 | sp\|P60900\|PSA6_HUMAN | component of the 20S core proteasome; proteolytic degradation |
| PSMA7 | sp\|O14818\|PSA7_HUMAN | component of the 20S core proteasome; proteolytic degradation |
| PSMB1 | sp\|P20618\|PSB1_HUMAN | non-catalytic component of 20S core proteasome complex |
| PSMB2 | sp\|P49721\|PSB2_HUMAN | non-catalytic component of 20S core proteasome complex |
| PSMB3 | sp\|P49720\|PSB3_HUMAN | non-catalytic component of 20S core proteasome complex |
| PSMB4 | sp\|P28070\|PSB4_HUMAN | non-catalytic component of 20S core proteasome complex |
| PSMB5 | sp\|P28074\|PSB5_HUMAN | component of the 20S core proteasome; proteolytic degradation |
| PSMB6 | sp\|P28072\|PSB6_HUMAN | one of the components of the 20S proteasome which is involved in the proteolytic degradation of many intracellular proteins |
| PSMB7 | sp\|Q99436\|PSB7_HUMAN | component of the 20S core proteasome; proteolytic degradation |
| PSMC1 | sp\|P62191\|PRS4_HUMAN | component of 26S proteasome |
| PSMC2 | sp\|P35998\|PRS7_HUMAN | component of 26S proteasome |
| PSMC3 | sp\|P17980\|PRS6A_HUMAN | component of 26S proteasome |
| PSMC4 | sp\|P43686\|PRS6B_HUMAN | component of 26S proteasome |
| PSMC5 | sp\|P62195\|PRS8_HUMAN | component of 26S proteasome |
| PSMC6 | sp\|P62333\|PRS10_HUMAN | component of 26S proteasome |
| PSMD1 | sp\|Q99460\|PSMD1_HUMAN | component of 26S proteasome |
| PSMD11 | sp\|O00231\|PSD11_HUMAN | component of 26S proteasome |
| PSMD12 | sp\|O00232\|PSD12_HUMAN | component of 26S proteasome |
| PSMD13 | sp\|Q9UNM6\|PSD13_HUMAN | component of 26S proteasome |
| PSMD14 | sp\|O00487\|PSDE_HUMAN | component of 26S proteasome |
| PSMD2 | sp\|Q13200\|PSMD2_HUMAN | component of 26S proteasome |
| PSMD3 | sp\|O43242\|PSMD3_HUMAN | component of 26S proteasome |
| PSMD4 | sp\|P55036\|PSMD4_HUMAN | component of 26S proteasome |
| PSMD6 | sp\|Q15008\|PSMD6_HUMAN | component of 26S proteasome |
| PSMD7 | sp\|P51665\|PSMD7_HUMAN | component of 26S proteasome |
| PSMD8 | sp\|P48556\|PSMD8_HUMAN | component of 26S proteasome |
| PSME4 | sp\|Q14997\|PSME4_HUMAN | component of proteasome involved in DNA repair; recognizes acetylated histones |
| **Chaperones** | | |
| BAG2 | sp\|O95816\|BAG2_HUMAN | co-chaperone |
| BAG6 | sp\|P46379\|BAG6_HUMAN | chaperone |
| CANX | sp\|P27824\|CALX_HUMAN | quality control protein in ER; interacts with newly synthesized glycoproteins in ER |
| CLGN | sp\|O14967\|CLGN_HUMAN | chaperone |
| CLPX | sp\|O76031\|CLPX_HUMAN | conditional chaperone |
| DNAJA2 | sp\|O60884\|DNJA2_HUMAN | co-chaperone |
| DNAJA3 | sp\|Q96EY1\|DNJA3_HUMAN | co-chaperone |
| DNAJB1 | sp\|P25685\|DNJB1_HUMAN | chaperone |
| DNAJB11 | sp\|Q9UBS4\|DJB11_HUMAN | co-chaperone |
| DNAJB12 | sp\|Q9NXW2\|DJB12_HUMAN | co-chaperone |
| DNAJB2 | sp\|P25686\|DNJB2_HUMAN | co-chaperone |
| DNAJB4 | sp\|Q9UDY4\|DNJB4_HUMAN | co-chaperone |
| DNAJB6 | sp\|O75190\|DNJB6_HUMAN | co-chaperone |
| DNAJC7 | sp\|Q99615\|DNJC7_HUMAN | co-chaperone |
| GET4 | sp\|Q7L5D6\|GET4_HUMAN | part of the cytosolic protein quality control complex (BAG6/BAT3 complex) |
| MDN1 | sp\|Q9NU22\|MDN1_HUMAN | chaperone |
| SDF2 | sp\|Q99470\|SDF2_HUMAN | co-chaperone |
| SDF2L1 | sp\|Q9HCN8\|SDF2L_HUMAN | co-chaperone |
| SGTA | sp\|O43765\|SGTA_HUMAN | co-chaperone |
| **Degradation Proteins** | | |
| ABCE1 | sp\|P61221\|ABCE1_HUMAN | hydrolase |
| AMFR | sp\|Q9UKV5\|AMFR_HUMAN | E3 ubiquitin ligase |
| ERLIN2 | sp\|O94905\|ERLN2_HUMAN | mediates ERAD |
| FAF2 | sp\|Q96CS3\|FAF2_HUMAN | ERAD |
| FBXO3 | sp\|Q9UK99\|FBX3_HUMAN | substrate recognition component of the SCF-type E3 ubiquitin ligase complex |
| HM13 | sp\|Q8TCT9\|HM13_HUMAN | protease |
| HUWE1 | sp\|Q7Z6Z7\|HUWE1_HUMAN | E3 ubiquitin ligase |
| ILVBL | sp\|A1L0T0\|ILVBL_HUMAN | lipid degradation; lyase in phytosphingosine degradation pathway |
| NBR1 | sp\|Q14596\|NBR1_HUMAN | ubiqutin binding autophagy |
| NPLOC4 | sp\|Q8TAT6\|NPL4_HUMAN | ubiquitin dependent degradation |
| PELO | sp\|Q9BRX2\|PELO_HUMAN | involved in the degradation of defective ribosomes through the activation of the No-Go Decay (NGD) pathway |
| PLAA | sp\|Q9Y263\|PLAP_HUMAN | protein ubiquitination, sorting, & degradation |
| RAD23B | sp\|P54727\|RD23B_HUMAN | receptor in proteasomal degradation |
| RNF126 | sp\|Q9BV68\|RN126_HUMAN | E3 ubiquitin ligase |
| SEL1L | sp\|Q9UBV2\|SE1L1_HUMAN | participates in the degradation of misfolded endoplasmic reticulum proteins |
| STUB1 | sp\|Q9UNE7\|CHIP_HUMAN | E3 ubiquitin ligase |
| SYVN1 | sp\|Q86TM6\|SYVN1_HUMAN | E3 ubiquitin ligase |
| UBAC2 | sp\|Q8NBM4\|UBAC2_HUMAN | negatively regulates Wnt signaling; Ubiquitin-associated domain-containing protein 2 |
| UBE3A | sp\|Q05086\|UBE3A_HUMAN | E3 ubiquitin ligase |
| UBL4A | sp\|P11441\|UBL4A_HUMAN | protein quality control; sorts proteins to the ER or to the proteasome for degradation |
| UBQLN1 | sp\|Q9UMX0\|UBQL1_HUMAN | regulator of a variety of protein degradation processes |
| UBQLN2 | sp\|Q9UHD9\|UBQL2_HUMAN | protein ubiquitination & degradation; ERAD |
| UFD1 | sp\|Q92890\|UFD1_HUMAN | ubiquitin dependent proteolytic pathway |
| USP11 | sp\|P51784\|UBP11_HUMAN | protease; deubiquitinating enzyme |
| USP19 | sp\|O94966\|UBP19_HUMAN | deubiquitinating enzyme |
| VCP | sp\|P55072\|TERA_HUMAN | DNA site repair; transport of ubiquitinated proteins |
| WDR48 | sp\|Q8TAF3\|WDR48_HUMAN | regulator of the deubiquitination pathway that activates USP1, USP12, USP 46 |
| WDTC1 | sp\|Q8N5D0\|WDTC1_HUMAN | substrate receptor for CUL4-DDB1 E3 ubiquitin-protein ligase complex |
| YME1L1 | sp\|Q96TA2\|YMEL1_HUMAN | ATP-dependent protease that catalyzes the degradation of folded and unfolded proteins |
| **Trafficking Proteins** | | |
| APOL2 | sp\|Q9BQE5\|APOL2_HUMAN | lipid transport |
| ARL1 | sp\|P40616\|ARL1_HUMAN | GTP binding protein that recruits effectors |
| ARL6IP5 | sp\|O75915\|PRAF3_HUMAN | regulates the taurine/glutamate concentrations & modulates glutamate transport |
| ATAD1 | sp\|Q8NBU5\|ATAD1_HUMAN | translocase |
| AUP1 | sp\|Q9Y679\|AUP1_HUMAN | translocation of terminally misfolded proteins from the ER lumen to the cytoplasm |
| CNIH4 | sp\|Q9P003\|CNIH4_HUMAN | involved in trafficking GPCRs from ER to cell surface |
| COPB1 | sp\|P53618\|COPB_HUMAN | coatomer for ER-Golgi transport |
| COPG2 | sp\|Q9UBF2\|COPG2_HUMAN | coatomer that binds to dilysine motifs and mediates protein transport in the ER -Golgi network |
| ERGIC1 | sp\|Q969X5\|ERGI1_HUMAN | involved in the ER-Golgi transport |
| ESYT2 | sp\|A0FGR8\|ESYT2_HUMAN | tethers ER to cell membrane & plays a role in FGF signaling; binds glycerophospholipids and may play a role in cellular lipid transport |
| EXOC4 | sp\|Q96A65\|EXOC4_HUMAN | component of the exocyst complex involved in the docking of exocytic vesicles; protein transport |
| FKBP8 | sp\|Q14318\|FKBP8_HUMAN | Peptidyl-prolyl cis-trans isomerase FKBP8 |
| IGF2R | sp\|P11717\|MPRI_HUMAN | cell trafficking receptor |
| KIFC2 | sp\|Q96AC6\|KIFC2_HUMAN | retrograde transport |
| KTN1 | sp\|Q86UP2\|KTN1_HUMAN | kinesin driven vesicle movement |
| MON2 | sp\|Q7Z3U7\|MON2_HUMAN | involved in membrane trafficking of cargo proteins |
| NUP210 | sp\|Q8TEM1\|PO210_HUMAN | acts as a nucleoporin which is essential for nuclear pore formation and structural integrity of the nuclear pores |
| NUP93 | sp\|Q8N1F7\|NUP93_HUMAN | important component of the nuclear pore complex (NPC) assembly |
| PEX1 | sp\|O43933\|PEX1_HUMAN | protein dislocase complex |
| RAB14 | sp\|P61106\|RAB14_HUMAN | acts as a GTPase which regulates intracellular membrane trafficking |
| RHOT1 | sp\|Q8IXI2\|MIRO1_HUMAN | involved in mitochondrial trafficking and its subcellular components by acting as a GTPase |
| RHOT2 | sp\|Q8IXI1\|MIRO2_HUMAN | involved in mitochondrial trafficking and its subcellular components by acting as a GTPase |
| SCFD1 | sp\|Q8WVM8\|SCFD1_HUMAN | involved in vesicular transport of proteins between Golgi Apparatus & Endoplasmic Reticulum |
| SLC27A4 | sp\|Q6P1M0\|S27A4_HUMAN | ligase & mediates transport of fatty acid |
| STX5 | sp\|Q13190\|STX5_HUMAN | mediates ER-Golgi transport |
| TCAF1 | sp\|Q9Y4C2\|TCAF1_HUMAN | positive regulator of the plasma membrane cation channel & recruits TRPM8 to cell surface |
| TMEM160 | sp\|Q9NX00\|TM160_HUMAN | transmembrane protein |
| TMEM201 | sp\|Q5SNT2\|TM201_HUMAN | nuclear movement during fibroblast polarization and migration |
| TMEM33 | sp\|P57088\|TMM33_HUMAN | positively regulates PERK and IRE1 signaling & immune signaling |
| TOMM70 | sp\|O94826\|TOM70_HUMAN | receptor in the translocase complex (TOM); required for mitochondrial transport |
| TOR1AIP1 | sp\|Q5JTV8\|TOIP1_HUMAN | induces ATPase activity & involved in nuclear lamina assembly |
| XPO1 | sp\|O14980\|XPO1_HUMAN | regulates export of proteins from nucleus to cytosol |
| **Modification Proteins** | | |
| AASDHPPT | sp\|Q9NRN7\|ADPPT_HUMAN | transferase that catalyzes post-translational modification of target proteins by phosphopanteeheine. |
| ADPGK | sp\|Q9BRR6\|ADPGK_HUMAN | kinase |
| ALDH3A2 | sp\|P51648\|AL3A2_HUMAN | aldehyde dehydrogenase that catalyzes oxidation of aldehydes to fatty acid |
| ALG1 | sp\|Q9BT22\|ALG1_HUMAN | transferase |
| ARAF | sp\|P10398\|ARAF_HUMAN | positive regulator for myogenic differentiation |
| CDKAL1 | sp\|Q5VV42\|CDKAL_HUMAN | methylthiotransferase |
| CHPF | sp\|Q8IZ52\|CHSS2_HUMAN | transferase |
| DCAF8 | sp\|Q5TAQ9\|DCAF8_HUMAN | possible substrate receptor for CUl4-DDB1 E3 ubiqutin ligase complex |
| DHCR24 | sp\|Q15392\|DHC24_HUMAN | oxidoreductase in cholesterol biosynthesis; involved in the reduction of sterol intermediates |
| DHRS7B | sp\|Q6IAN0\|DRS7B_HUMAN | putative oxidoreductase |
| FDFT1 | sp\|P37268\|FDFT_HUMAN | squalene synthesis |
| IRAK1 | sp\|P51617\|IRAK1_HUMAN | protein kinase for S & T |
| MAGT1 | sp\|Q9H0U3\|MAGT1_HUMAN | transferase |
| MOGS | sp\|Q13724\|MOGS_HUMAN | involved in N-gluycan degradation; cleaves distal glucose residue from precursor |
| NAT14 | sp\|Q8WUY8\|NAT14_HUMAN | probable acetyltransferase |
| NEU1 | sp\|Q99519\|NEUR1_HUMAN | catalyzes the removal of sialic acid from glycoproteins and glycolipids |
| P3H1 | sp\|Q32P28\|P3H1_HUMAN | prolyl 3-hydroxylase |
| PFKM | sp\|P08237\|PFKAM_HUMAN | phosphorylates D-fructose 6-phosphate |
| PGM3 | sp\|O95394\|AGM1_HUMAN | catalyzes conversion during the synthesis of UDP-GlcNAc |
| PON2 | sp\|Q15165\|PON2_HUMAN | hydrolyzes lactones (cyclic esters) and other aromatic esters |
| PPP2R5C | sp\|Q13362\|2A5G_HUMAN | phosphatase activity |
| PPP6R1 | sp\|Q9UPN7\|PP6R1_HUMAN | subunit component of the regulatory protein phosphatase 6 (PP6) |
| PRKAR1A | sp\|P10644\|KAP0_HUMAN | regulatory component of cAMP signaling by acting as a subunit of cAMP dependent protein kinases |
| PRKAR1B | sp\|P31321\|KAP1_HUMAN | regulatory subunit of cAMP protein kinases in the cAMP signal pathway |
| QPCTL | sp\|Q9NXS2\|QPCTL_HUMAN | involved in the biosynthesis of pyroglutamyl peptides |
| RAF1 | sp\|P04049\|RAF1_HUMAN | phosphorylates serine and threonine |
| RANBP2 | sp\|P49792\|RBP2_HUMAN | facilitates the conjugation of two proteins SUMO1 & SUMO2 by acting as a ligase |
| TMX3 | sp\|Q96JJ7\|TMX3_HUMAN | Participates in the folding of proteins containing disulfide bonds |
| UGGT1 | sp\|Q9NYU2\|UGGG1_HUMAN | reglucosylates proteins |
| UGGT2 | sp\|Q9NYU1\|UGGG2_HUMAN | reglucosylates proteins |
| **Uncategorized Proteins** | | |
| ACSL4 | sp\|O60488\|ACSL4_HUMAN | ligase that catalyzes the conversion of long-chain fatty acids to acyl-CoA; lipid metabolism |
| APMAP | sp\|Q9HDC9\|APMAP_HUMAN | strong arylesterase activity with beta-naphthyl acetate and phenyl acetate |
| ATRIP | sp\|Q8WXE1\|ATRIP_HUMAN | involved in DNA repair and damage respons |
| ATXN10 | sp\|Q9UBB4\|ATX10_HUMAN | induces neuritogenesis by activating the Ras-MAP kinase pathway |
| CLPTM1 | sp\|O96005\|CLPT1_HUMAN | binds/traps GABAA receptors in the ER; modulates GABAergic transmission |
| EIF2B3 | sp\|Q9NR50\|EI2BG_HUMAN | translation initiation factor; GEF-like activity |
| FANCI | sp\|Q9NVI1\|FANCI_HUMAN | DSB repair via HR; critical for interstrand crosslink repair; promotes Mono ubiquitination of FANCD2 |
| ITGA6 | sp\|P23229\|ITA6_HUMAN | receptor for laminin on platelets; essential for NRG1-ERBB signaliing |
| ITM2B | sp\|Q9Y287\|ITM2B_HUMAN | processes amyloid-beta A4 precursor protein (APP) & acts as an inhibitor of amyloid-beta peptide aggregation |
| MLF2 | sp\|Q15773\|MLF2_HUMAN | myeloid leukemia factor |
| NDUFS3 | sp\|O75489\|NDUS3_HUMAN | subunit of NADH Dehydrogenase (Complex I) of the Electron Transport Chain |
| RMDN3 | sp\|Q96TC7\|RMD3_HUMAN | induces apoptosis when overexpressed (may be involved in the mitochondria/cytochrome C apoptosis pathway; calcium homeostasis |
| SPTLC1 | sp\|O15269\|SPTC1_HUMAN | a component in the serine palmitoyltransferase multisubunit enzyme (SPT) |
| TRAFD1 | sp\|O14545\|TRAD1_HUMAN | Negative feedback regulator; regulates toll-like receptor 4 |
| ZC3H18 | sp\|Q86VM9\|ZCH18_HUMAN | Enables mRNA cap binding complex binding activity and protein-macromolecule adaptor activity |
